# Delayed benefits for fallow bucks: more fights decrease same day mating success, but may increase matings the next day

**DOI:** 10.1101/2022.05.19.492486

**Authors:** Alessandra Bateman-Neubert, Elodie F. Briefer, Alan G. McElligott

## Abstract

Dominance hierarchies help to reduce unnecessary fights and associated costs during the mating season. Fallow deer (*Dama dama*) typically have high levels of male-male competition and strong reproductive skew. Nevertheless, how male dominance and daily fight rates affect mating success remains unknown. We used a two-year dataset from a large population of tagged fallow deer (620-689 individuals), to calculate male dominance ranks based on their agonistic interactions prior to the mating season (‘prerut’), in order to then examine how rank is related to fight rates and mating success during the mating season (‘rut’). Overall, higher-ranked males fought at least twice a day on a higher proportion of days during the rut and secured more matings. Males engaging in more than 10 fights per day were less likely to secure a mating that same day, and those males exceeding 15 fights per day secured no matings at all. Nevertheless, males with the highest numbers of fights (i.e. 15-21 fights per day) on a given day had higher mating success on subsequent days. Although higher-ranked males secured most matings during the rut, their fight rates decreased towards the end. We propose that engaging in more fights negatively affects daily individual mating success, but may benefit mating success on subsequent days, and potentially increase long-term fitness benefits. Additionally, engaging in more fights as the rut progresses probably allows lower-ranked males to secure some matings before the availability of oestrous females ends for almost a year.

**SIGNIFICANCE STATEMENT:** Fighting carries a risk of injury and high energetic costs. Male fallow deer establish dominance hierarchies, that help reduce unnecessary fights among individuals of different competitive abilities. However, whether high-ranked males fight more or less is yet unknown. By calculating social ranks of fallow bucks before the start of their mating period, we show that males of higher social status do fight and mate more during the mating season (rut). Furthermore, by investigating how investment in fights affects individual mating success that same day and the next day, we find that males that fight more cause a decrease in their immediate daily mating success, but can potentially increase their chances of mating in subsequent days. Thus, to fight more may allow males to climb the hierarchical social ladder, hence increasing longer-term fitness benefits associated to higher ranks.

## INTRODUCTION

In species with polygynous mating systems, dominance status depends on male competitive ability, and impacts on female mate choice and reproductive success (Fanjul & Zenuto, 2017; Farrell et al., 2011; Willisch et al., 2012; Wyman et al., 2021). High degrees of polygyny are characterized by strong reproductive skew, where a few males get most copulations during a mating season (Emlen & Oring, 1977). Dominance position within a social hierarchy determines individual mating success, where dominance can be achieved through a variety of aggressive (e.g. fights) and non-aggressive (e.g. avoidance, grooming) behaviors (Ellis, 1995; Lewis, 2022).

Direct intrasexual competition (e.g. fighting) between males is an important factor for establishing certainty on dominance relationships and for gaining access to females (Clutton-Brock & Huchard, 2013; Silk et al., 2019). However, fights carry high risks of injury and great energetic costs (Emery-Thompson & Georgiev, 2014; Lukas & Clutton-Brock, 2014). To reduce such costs, social dominance hierarchies are commonly established and function to limit unnecessary conflict, namely when lower quality individuals avoid fighting with more dominant males (Lewis, 2022; Magaña et al., 2011; Willisch & Neuhaus, 2010). Increased knowledge of their social environment may allow individuals to optimize the directionality of aggressive behaviors (i.e. who to fight with according to social rank; Hobson et al., 2021). Moreover, males may use information on social environment to modulate the intensity of aggressive behaviors, where costlier behaviors are invested in fighting closely-ranked males, when winning results in greater benefits (Dehnen et al., 2022b).

In polygynous ungulates, males use different mating tactics depending on their quality, rank and specific environmental circumstances in order to increase access to females and mating success (Bowyer et al., 2020). Mating tactics can differ in ‘ways’ to approach a female or ‘times’ when to approach a female. Mating tactics used by dominant males tend to be more effective, while subordinates are likely to use alternative mating tactics in order to secure a mating while avoiding contact with higher-ranked males (Foley et al., 2018; McElligott et al., 2001; McElligott & Hayden, 2000). As the mating season advances, body condition of males decreases and information on these changes in body condition is communicated to the rest of the population through, for example, changes in the structure of their vocalizations (McElligott et al., 2003; Mysterud et al., 2008; Vannoni & McElligott, 2009; Wyman et al., 2021). Furthermore, at the end of the mating season, dominant males are likely to decrease investment in mating-related activities and switch to self-maintenance behaviors (e.g. feeding, resting). As a result, subordinates have increased likelihood of securing a mating at that time (Farrell et al., 2011).

The fallow deer (*Dama dama*) is a polygynous species with strong reproductive skew, in which males actively compete for access to females (Clutton-Brock et al., 1988; McElligott & Hayden, 2000). Fallow bucks can be described as socially immature (≤ 3 years old) or mature (≥ 4 years old), according to the behaviors they exhibit during the mating season, also called ‘rut’ (McElligott et al., 1999). Socially immature males rarely participate in contests during the rut, avoiding fighting costs and potential injuries, unlike socially mature bucks, that actively participate (McElligott et al., 1998). Socially mature males shape their dominant-subordinate relationships before the mating season or ‘prerut’, mainly through low-cost, non-contact, agonistic interactions. Dominance-relationships among fallow deer males may slightly shift throughout the mating season, as is the case for other species with hierarchical structures (Chase et al., 2022; Shizuka & McDonald, 2012). However, the main resulting ranks from prerut interactions tend to persist during the breeding period (Ciuti & Apollonio, 2016; McElligott et al., 1998).

Investment in fights increases as a function of male individual condition and number of available females in both fallow deer and red deer (*Cervus elaphus*), (McElligott et al., 1998, 2003; Mysterud et al., 2008). Dominant fallow deer males, usually in prime-age (6-7 years old), are of greater body mass and size and tend to have the highest mating success (McElligott et al., 2001). However, whether the dominance rank established before the mating season affects the number of fights during the rut in male fallow deer, remains unknown. Moreover, changes in body condition throughout the rut can affect individual investment in fights and matings, leading to possible changes in the relationship between male rank, fight rate and mating success.

We investigated the relationship between male fallow deer prerut social rank and their fight rate, fighting success and mating success and the relationship between the number of fights in a day and mating success. We further investigated whether there is a rank-dependent change in the fight rate and mating success of males throughout the rut, in order to understand the costs and benefits of fighting. We predicted that higher-ranked males would have lower fight rates and higher mating success at the peak of the rut, given that these individuals are of higher quality and are potentially avoided by lower-ranked males (Willisch & Neuhaus, 2010). We also anticipated that males with highest numbers of fights per day would have lower mating success, linked to a potential trade-off in energy and time investment between fighting *versus* mating. However, we predicted that males that more often win fights would gain more matings, given that winning is an attractive trait to females (Gerber et al., 2010; House et al., 2019). Lastly, we expected males to change their investment in fights and matings as the rut progressed, given the energetically costly rut-related activities and the existence of various mating tactics expressed in this species (Ciuti & Apollonio, 2016)

## METHODS

### Study site and population

Data used for this study were collected over two years (1994-1995), as part of a long-term study of European fallow deer (*Dama dama*) population in Phoenix Park (53°22′ N, 6°21′ W), Dublin, Ireland. The park is approximately 709 ha, of which 80% is pasture and 20% is woodland, providing enough foraging resources for the herd to survive without additional food supply. Deer identities in this study were known as a result of the tagging efforts by the park authorities (McElligott & Hayden, 2000). Population size was of 620 deer in 1994 (315 females and 200 males), and 689 deer in 1995 (353 females and 200 males) (McElligott & Hayden, 2000). Only socially mature males seen interacting with each other were included for analyses in this study (Table 1).

**Table 1.**
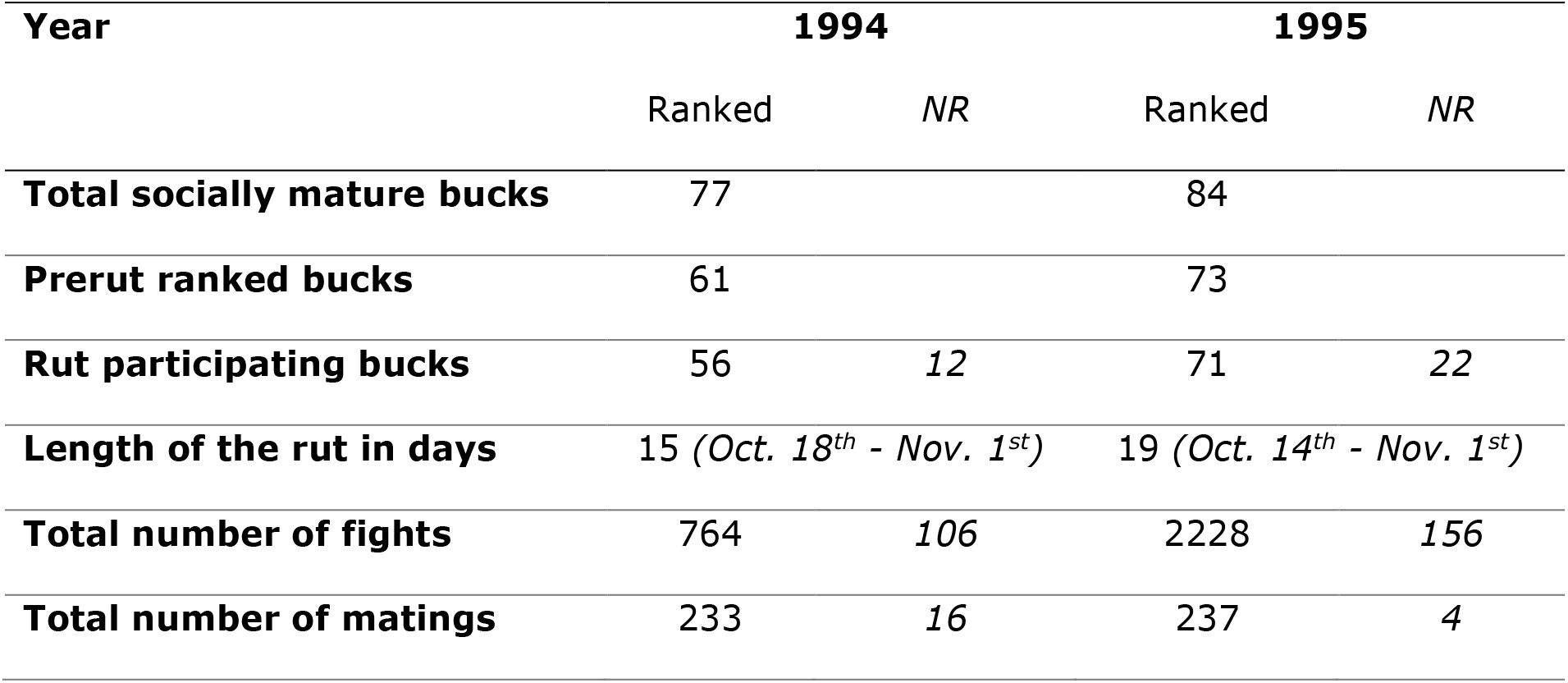
Prerut fallow bucks, participating ranked bucks and non-ranked bucks during the rut and total number of fights and matings for 1994 and 1995. NR stands for non-ranked bucks

### Data collection

Data collection was done by a group of 6-11 observers during the mating season of each year. The season is divided in two periods; the prerut starts when the last male has cleaned the velvet from their antlers and finishes the day before the first mating of the season occurs and the rut starts with the first mating of the season and ends with the last mating. The prerut started at the end of August, with a total of 32 days sampled in 1994 and 39 days in 1995. The rut lasted 15 days in 1994 and 19 days in 1995, starting mid-October and ending on the 1^st^ of November, both years (McElligott & Hayden, 2000) (Table 1).

### David’s Score method

The outcomes of agonistic interactions between male dyads are categorised as win/loss, where one male wins (i.e. chases off opponent) and the other loses (i.e. retreats from fight location) that interaction, or as undetermined, where none clearly wins nor loses. A win or a loss has consequences on male’s subsequent agonistic behavior, known as the winner-loser effect, enhancing dominant or subordinate behaviors (Hsu & Wolf, 2001; Oldham et al., 2020). Clear outcomes of agonistic interactions between males during the prerut were used to calculate the dominance rank based on the David’s Score (DS) method. The DS is a suitable ranking method when large numbers of interactions have been observed, as it considers repeated interactions between the parties of a dyad in which win/loss asymmetries may be clarified. This method avoids an artificially increased or lowered position of the dominance rank of certain individuals due to potential minor deviations from the general dominance direction or simple observer mistakes. Nevertheless, the ranking position of an individual may be erroneously increased or lowered, when this individual interacts with a small proportion of the population only, or when it obtains only wins or only losses in all its interactions. In this case, it would be appropriate to exclude it from the dominance rank calculation (Gammell et al., 2003).

To calculate the dominance hierarchy, mature male dyads were selected from the prerut period. Only agonistic interactions with a clear winner (i.e. interactions with resolved outcomes) were selected, considering that the method uses dyadic interactions where one of the parties ‘scores’ while the other ‘loses’ (David, 1987; Gammell et al., 2003). First, the percentage of interactions that each buck had with other mature males during the prerut was calculated. Those males interacting with less than 10% of the total mature male population were removed, which included those males that had only wins or only losses (Briefer et al., 2010; Gammell et al., 2003). Scores were transformed into a hierarchical sequence (1-n), where number 1 determined the highest score, hence the top dominant male that year, followed by males with lower scores until the last male in the hierarchy that year (‘n’) with the lowest score, 61 in 1994 and 73 in 1995 (Table 1).

### Statistical analyses

All data analyses and graphical representations were performed in R software version 3.6.1 with RStudio (an integrated development environment for R). Linear Mixed-effects Models (LMM’s) (lme4 library; Bates et al., 2015) were used to investigate the following effects:

1) the effect of rank on the number of fights per buck per day, on the outcome of those fights, and on the number of matings per buck per day (in three separate models); 2) the effect of fights on the number of matings per buck per day, and on the number of matings per buck the next day (in two separate models); and 3) the effect of the outcome of fights on the number of matings per buck per day, and on the number of matings per buck the next day (in two separate models).

To test the assumptions of normality and homoscedasticity, residuals of the models were visualized in Q-Q plots and scatterplots using the function “simulateResiduals” (DHARMa library; Hartig, 2020). When the assumptions were not met, a quadratic, log or binary transformation of the response variable was performed. All response variables (fights per buck per day, matings per buck per day, matings per buck the next day and outcome of fights) had a better fit when binary-transformed and were thus input in Generalized Linear Mixed-effect Models (GLMM) using the glmer function (lme4 library; Bates et al., 2015), with binary family, instead of LMM. To this aim, data were transformed as follows; the number of fights per buck per day as: > 1 fight = 1 and 0-1 fight = 0, the number of matings per buck per day and the number of matings per buck the next day as: > 0 matings = 1 and 0 matings = 0, and the outcome of fights as: win = 1 and loss =0. Thus, fight rate should be interpreted in our results as the proportion of days during which a buck fought at least twice (i.e. ≥ 2 fights per buck per day) or less (i.e. 0-1 fights), outcome of fights as the proportion of days that males won or lost fights (i.e. won or lost ≥ 1 fights per buck per day), and mating rate as the proportion of days during which a buck mated (i.e. ≥ 1 mating per buck per day) or not at all (i.e. 0 matings).

For each model we decided if fixed effects (rank, fights per buck per day, outcome) would be input as linear, quadratic or logarithmic terms based on the model’s Akaike Information Criterion adjusted for small sample sizes (AICc) (Burnham et al., 2011). The model with the smallest AICc values was chosen, and the fixed effects were fit as linear only, linear and quadratic, or logarithmic terms accordingly. If in one case, the AICc values of two or three models differed by less than 2 (Delta AICc: DAICc < 2), both models were considered to have support and thus were considered as potential fits to explain the data. Nevertheless, if these similarly supported models (DAICc < 2) differed in the number of parameters used (K), the one with the lower K was chosen as the most parsimonious answer (Burnham et al., 2011). Only the selected models are described in the result section; for further information on model selection, see Supplementary Tables S1-S5.

All models included buck identity, itself crossed with buck identity nested within year, to control for repeated measures of the same individual between and within years. The day of rut was added as an additional linear and quadratic fixed term to control for the daily change in fights and matings over the rut. The p-value for each fixed effect was obtained from a bootstrap test, based on 1000 replicates, through the comparison of a model with and a model without the fixed factor of interest (pbkrtest library; Halekoh & Højsgaard, 2014). However, when the bootstrap method failed, we used a Likelihood-Ratio Test instead.

Non-ranked males (n = 34) were not included in the analyses. To test the effect of fights on the number of matings per buck per day and on the next day, entries in which bucks had 0 fights and ≥ 1 mating/s the same day (n = 15) were excluded from the model for the following reason; as bucks that had 0 fights and 0 matings in one day were not recorded (i.e. during field data sampling, bucks that do not fight or mate are not registered), such calculations give a wrong output suggesting that 100% of bucks that did not fight obtained a mating otherwise.

Since the effect of the day of rut was significant (see Results), we further tested the effect of rank on fights and matings separately for the start, peak and end of the rut. The ruts of both years were divided into three independent periods, based on the mating activity. To determine the peak of the season, the day with the highest percentage of matings was selected for each year (15.45% of matings on days 7 and 10 of the rut of 1994; 13.5% on day 12 in 1995); those days with a percentage of matings of at least half of the maximum, were included in the ‘peak’ of the rut (days 6-11 with ≥ 7.7% of matings in 1994; days 9-15 with ≥ 6.8% in 1995; Supplementary Fig. S1). The days before the peak of the rut were attributed to the ‘start’ of the rut and the days after the peak, to the ‘end’ of the rut. In 1995, although day 17 had more than 6.8% of the matings, it was not included in the peak due to the discontinuity in the number of matings, as day 16 did not reach the minimum percentage (Supplementary Fig. S1). The three periods were then analyzed independently following the same procedure as described above. These new models included the same fixed and random effects as described above.

## RESULTS

### Group size and data distribution

Of the 134 ranked males, 128 males participated and had a total of 470 matings and 2992 fights over the ruts of two years. From the total of sampled observations, 95% of instances were males that had 0-6 fights per day, while only 5% of observations were bucks engaging in ≥ 7 fights in a day. The highest number of fights per day a buck was involved in was 21 fights and the highest number of matings secured by a buck during a day was 22 matings. The top 10 males secured 81% of the matings and were involved in 25% of the total fights. Additionally, 95 males (71%) that engaged in fights during the rut secured no matings at all (Table 2).

**Table 2.**
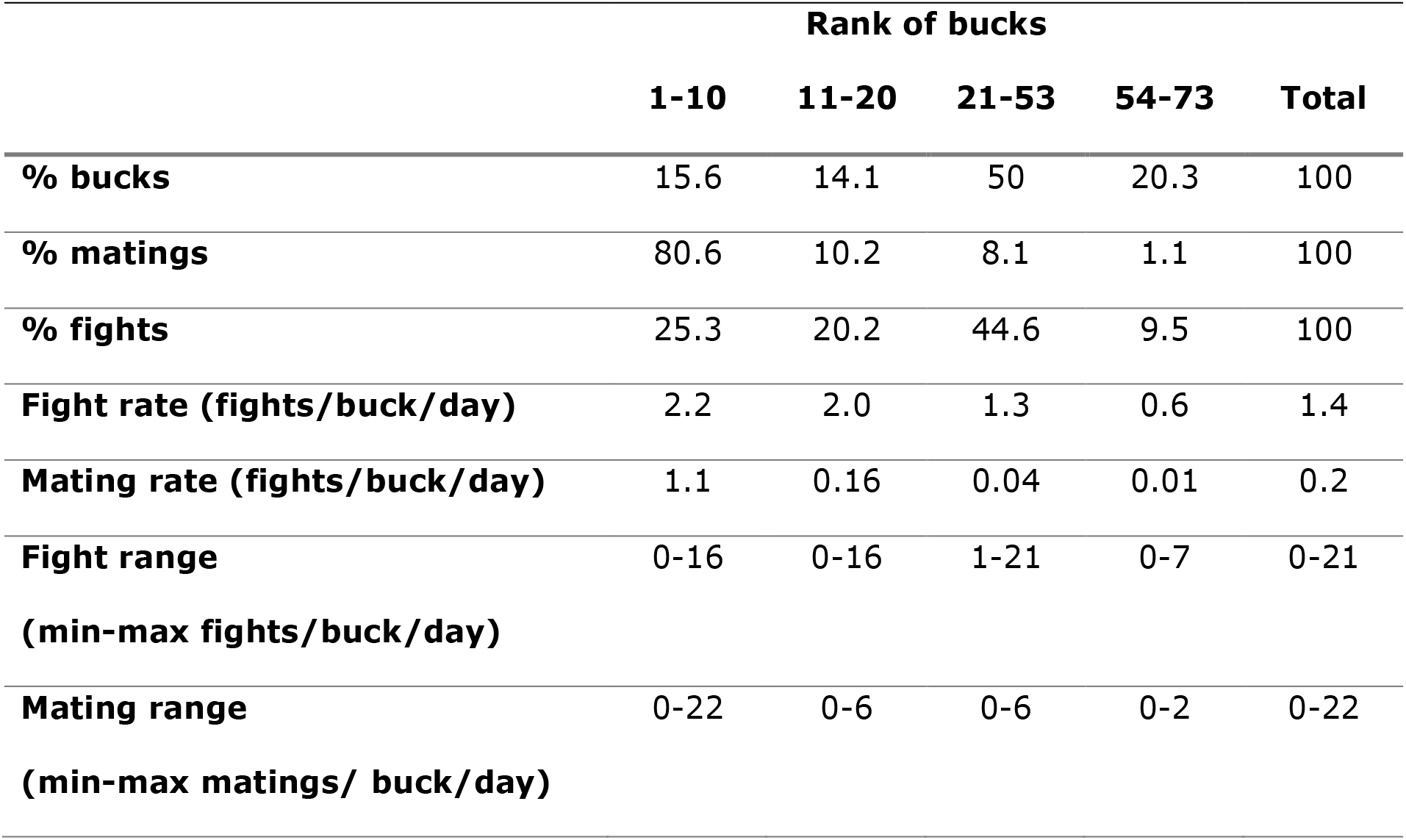
Results summary overview

### The effect of dominance, fight rate and fighting success

Higher-ranked males fought at least twice (i.e. ≥ 2 fights) per day on a higher proportion of days (calculated per buck per day; GLMM: quadratic relationship, p-value = 0.03; Fig. 1a) and won fights on a higher proportion of days (GLMM: quadratic relationship, p-value = 0.001; Fig. 1b) than lower-ranked males. This quadratic relationship between prerut rank and fighting success revealed by the model selection based on AICc (Supplementary Table S1), suggests that fighting success increases between mid-ranked and lower-ranked males. Higher-ranked males mated at least once (i.e. ≥ 1 mating) per day on a higher proportion of days during the rut compared to the lower-ranked ones (GLMM: logarithmic relationship, p-value = 0.001; Fig. 1c). The proportion of males that secured at least one mating was positively related to the number of fights they engaged in on that same day, up to a threshold (approximately 10 fights/buck/day), after what mating success decreased. Males with more than 15 fights a given day, secured 0 matings that same day (GLMM: quadratic relationship, p-value = 0.001; *Fig. 2a*). Instead, a higher proportion of males involved in the highest numbers of fights a given day, mated at least once on the next day, compared to those males engaging in fewer fights (GLMM: logarithmic relationship, p-value = 0.001; Fig. 2b). Finally, males that won a fight during a given day did not have greater mating success that same day (GLMM: p-value = 0.23; Fig. 3a), or the day after (GLMM: p-value = 0.20; Fig. 3b), than males that lost (For further information on model selection, see Supplementary Tables S1, S2 and S3).

**Fig. 1.**
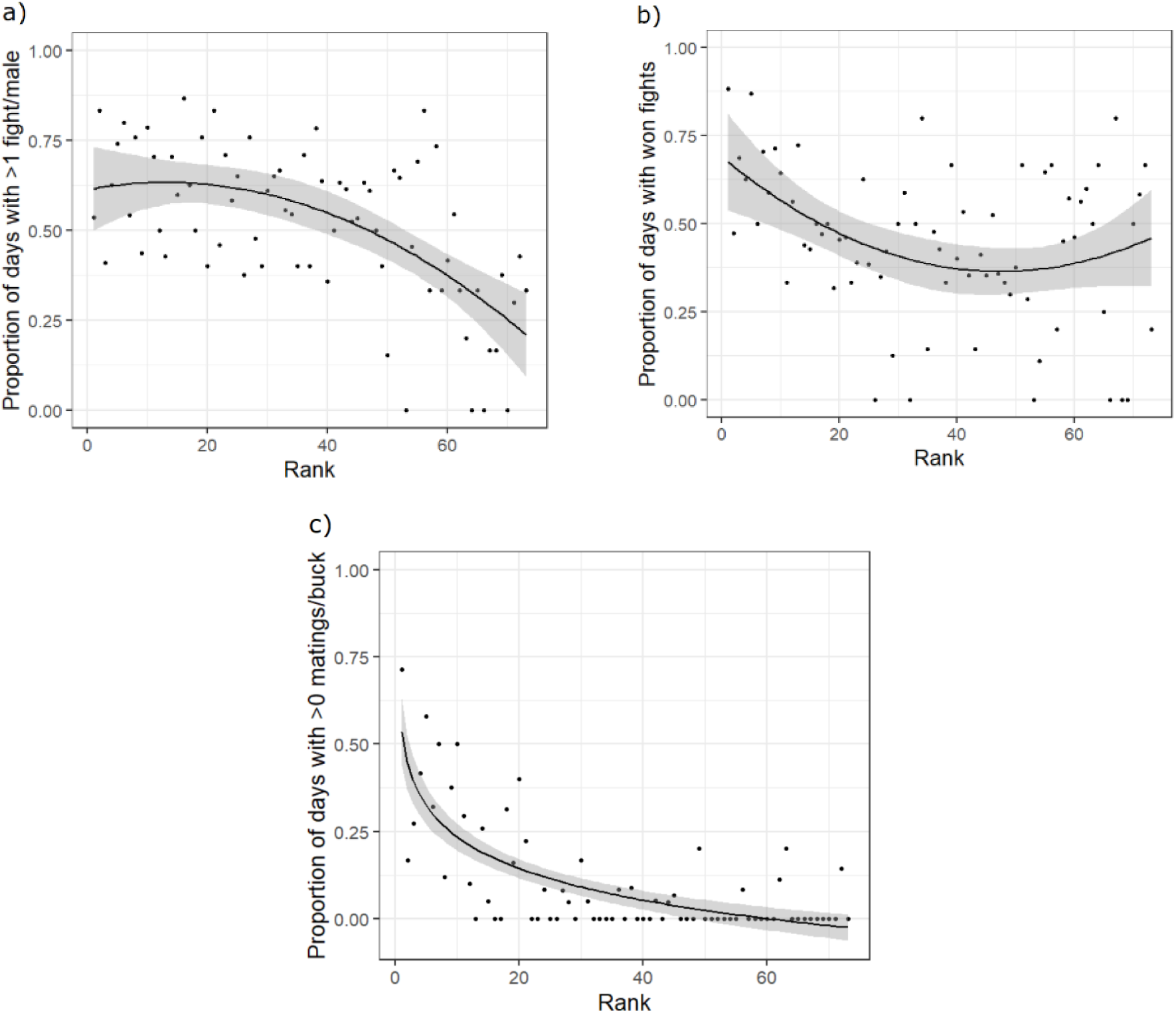
(a) Relationship (quadratic) between rank (prerut hierarchical dominance) and proportion of days that males fight at least twice (calculated per buck per day, binary transformed). (b) Relationship (quadratic) between rank (prerut hierarchical dominance) and proportion of days that males won fights (1 = win; 0 = loss). (c) Relationship (logarithmic) between rank (prerut hierarchical dominance) and proportion of days that males mate (calculated per buck per day, binary transformed). The black line represents the regression line and the grey areas indicate the 95% confidence interval

**Fig. 2.**
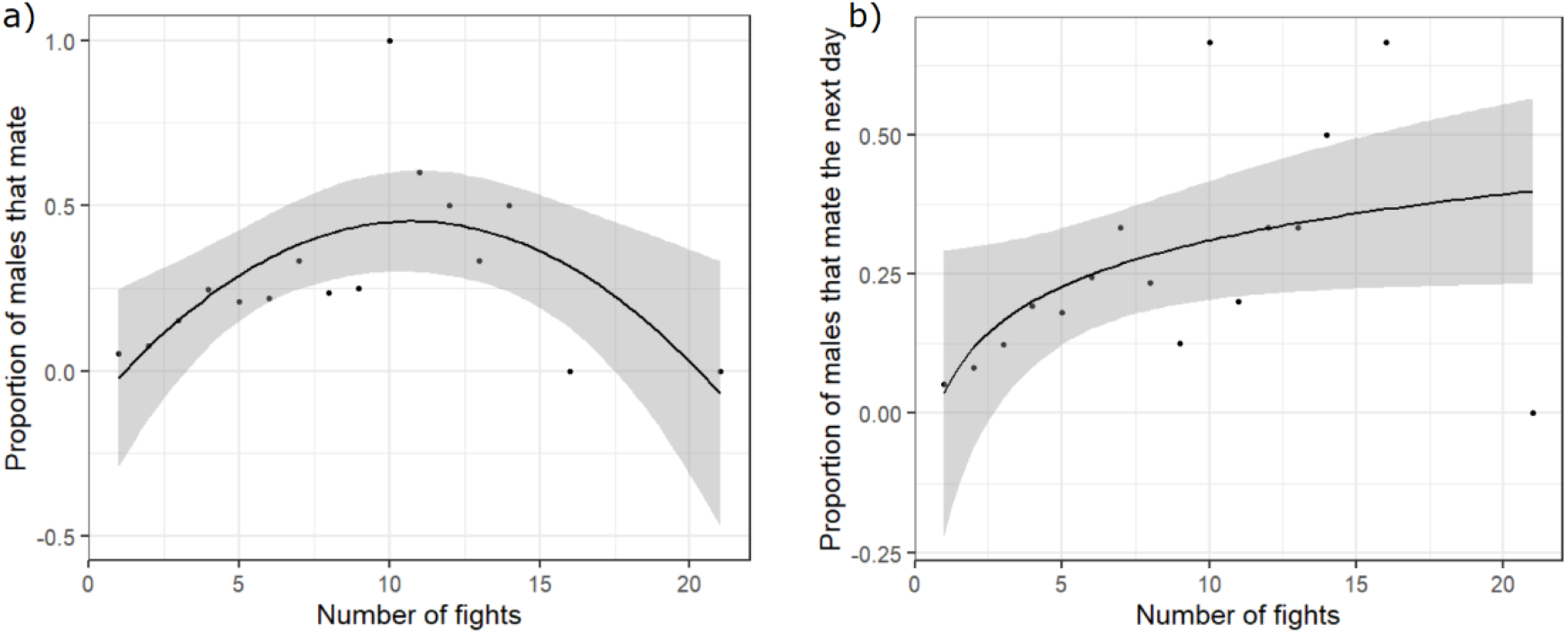
(a) Relationship (quadratic) between the number of fights (per buck per day) and proportion of males that mate at least once that same day (calculated per buck per day, binary transformed). (b) Relationship (logarithmic) between the number of fights (per buck per day) and proportion of males that mate at least once the next day (calculated per buck per day, binary transformed). The black line represents the regression line and the grey areas indicate the 95% confidence interval

**Fig. 3.**
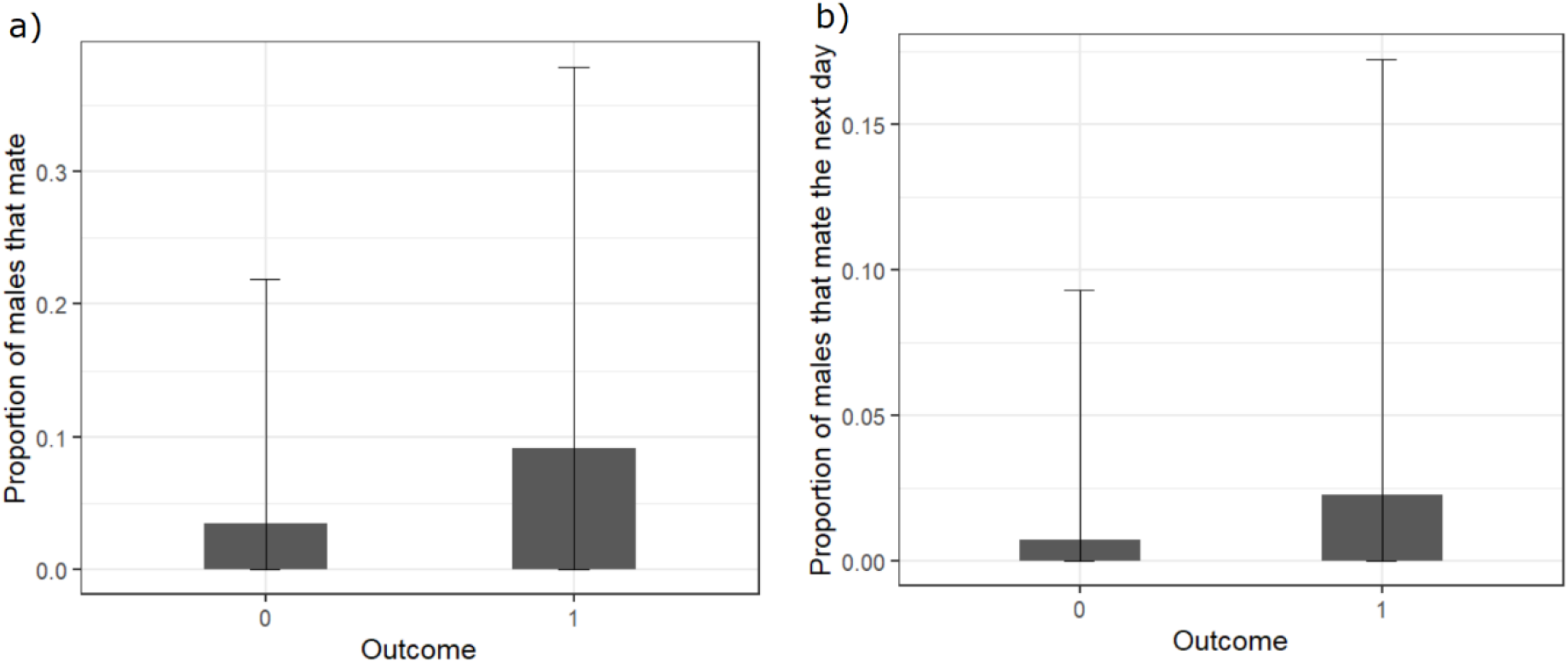
(a) Relationship between outcome of fights (1 = win; 0 = loss) and proportion of males that mate at least once that same day (calculated per buck per day, binary transformed). (b) Relationship between outcome of fights (1 = win; 0 = loss) and proportion of males that mate at least once the next day (calculated per buck per day, binary transformed). Error bars represent standard deviation. Both relationships were found to be not significant

### Differences between the three periods of the rut

Higher-ranked males fought at least twice per day on a higher proportion of days than lower-ranked males during the start (GLMM: quadratic relationship, p-value = 0.02; Fig. 4a) and peak (GLMM: linear relationship, p-value = 0.001; Fig. 4b) of the rut, but not at the end of the rut (GLMM: linear relationship, p-value = 0.11; Fig. 4c). In addition, higher-ranked males mated at least once per day on a higher proportion of days than lower-ranked males at the start (GLMM: logarithmic relationship, p-value = 0.001; Fig. 5a), peak (GLMM: logarithmic relationship, p-value = 0.001; Fig. 5b) and end (GLMM: logarithmic relationship, Chi-sq = 7.69, p-value = 0.01; Fig. 5c) of the rut. Higher-ranked males seemed to secure at least one mating more often at the start of the rut (Fig. 4a), while lower-ranked males did not mate nor fight during that period (Figs. 4a, 5a). Most males seemed to fight at least twice per day more often during the peak of the rut (Fig. 4b), while higher-ranked males still secured at least one mating more often than the rest of males (Fig. 5b). At the end of the rut, higher-ranked males seemed to secure at least one mating per day less often while lower-ranked males secured a few matings (Figs. 4c, 5c) (for further information on model selection, see Supplementary Tables S4 and S5).

**Fig. 4.**
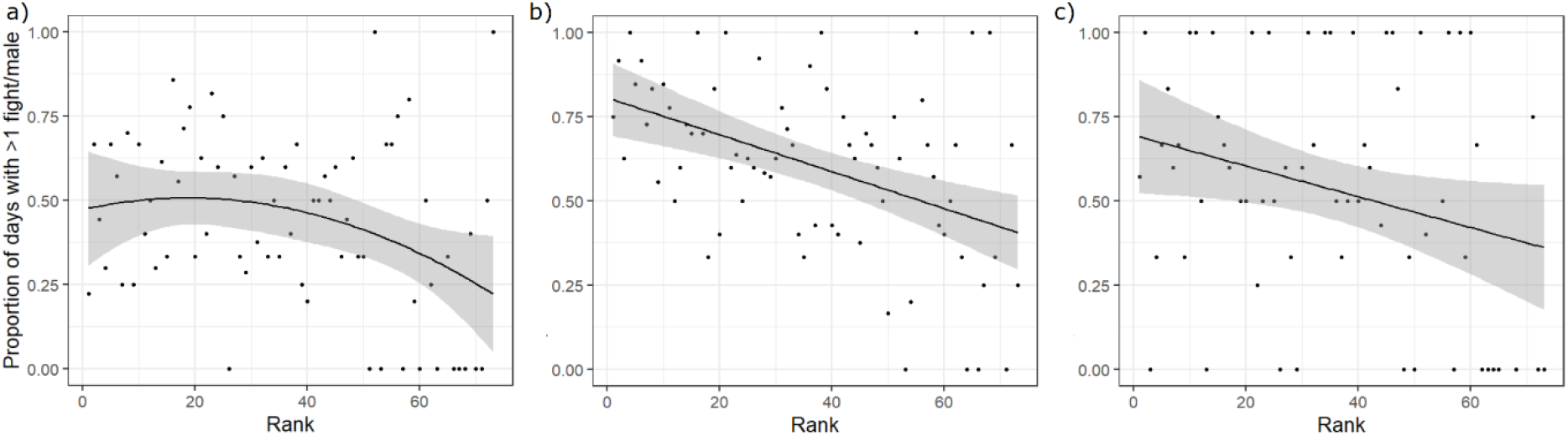
Relationship between rank (prerut hierarchical dominance) and proportion of days that males fight at least twice (calculated per buck per day, binary transformed) at (a) start (quadratic relationship), (b) peak (linear) and (c) end (linear) of the rut. The black line represents the regression line and the grey areas indicate the 95% confidence interval

**Fig. 5.**
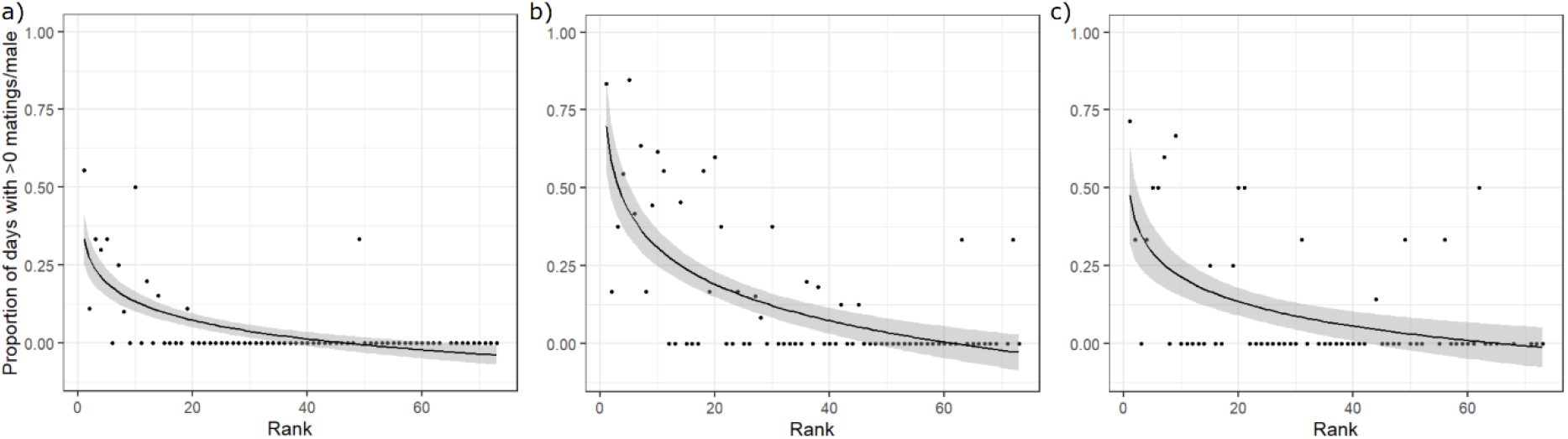
Relationship (logarithmic) between rank (prerut hierarchical dominance) and proportion of days that males mate (calculated per buck per day, binary transformed) at (a) start, (b) peak and (c) end of the rut. The black line represents the regression line and the grey areas indicate the 95% confidence interval

## DISCUSSION

We studied the relationship between dominance rank and the proportion of days during the rut on which socially mature fallow deer males were involved in at least two fights, and the effect of these two factors on their mating success (i.e. proportion of days that males mated at least once). We found that higher-ranked males fought at least twice a day on a higher proportion of days during the rut, won more fights and had higher mating success compared to lower-ranked males. Males engaging in more than 10 fights per day were less likely to secure a mating that same day, and those males exceeding 15 fights per day secured no matings at all. Nevertheless, males with the highest numbers of fights (i.e. 15-21 fights per day) on a given day had higher mating success on subsequent days, suggesting that males gain delayed longer-term benefits from fighting. Contrary to what was initially expected, we found no relationship between fighting success and mating success. Moreover, higher-ranked males fought at least twice per day on a lower proportion of days towards the end of the rut, compared to the start and peak of the rut. While changes in the proportion of days that males engage in at least two fights do not seem to affect mating success of higher-ranked males, it may allow for some lower-ranked males to secure a mating. Overall, our results suggest that males likely increase current costs by fighting more, in order to increase future short-term mating success (Dehnen et al., 2022a; Lewis, 2022; Tibbetts et al., 2022).

Our results indicate that higher-ranked males had at least two fights per day on a higher proportion of days throughout the rut compared to lower-ranked males. According to previous studies, higher-ranked fallow deer males, which are of higher quality, can afford to fight more while strongly reducing food intake and have an earlier onset of vocalizations (Apollonio & Di Vittorio, 2004; Vannoni & McElligott, 2009). Indeed, high-energy expenditure by successful males does not significantly affect their overwinter survival or lifetime success, besides the inherent risk of injury when fighting (Ciuti & Apollonio, 2016; McElligott et al., 2002, 2003). Moreover, experienced males assess their competitors for longer, with non-contact behaviors (e.g. parallel walk), before engaging in a fight, hence reducing costs of unnecessary fights (McElligott et al., 1998). Thus, higher-ranked males may engage in more fights, but their greater quality and experience may help reduce costs of fighting.

Our results indicate that a high number of fights per day negatively affects male mating success. The proportion of males that mated at least once increased with their number of fights, but only up to approximately 10 fights per day, after what fewer males secured at least one mating on that same day. Males engaging in the highest numbers of fights secured no matings that same day. How high aggression rates among males may affect their individual mating success may be related to females leaving areas of high conflict, as previously observed in fallow deer, or to time or energy budget constraints as observed in other species, e.g. California Sea Lions (*Zalophus californianus*), (Apollonio et al., 1989; Gerber et al., 2010; Naulty et al. 2013). However, 95% of males in our study did not exceed six fights per day, while only few bucks fought more than 10 fights in a day. Therefore, we suggest that higher-ranked males are likely to engage in few fights per day, while fighting during most rut days, hence maintaining their ranking position throughout the rut as a result.

Males that fought more on a given day appeared to have greater chances of securing at least one mating the following day compared to those males engaging in fewer fights. How engaging in high numbers of fights is related to high individual mating success the next day may potentially be associated to female choice. Indeed, females in lekking fallow deer populations choose males by preferred locations, indicating males that are better qualified to maintain that spot during the rut (Apollonio et al., 1989). Additionally, male mating success in fallow deer and red deer (*Cervus elaphus L*.) is related to male fighting success (Clutton-Brock et al., 1979, 1988; Moore et al., 1995). However, our results showed no relationship between fighting success and mating success. Thus, male mating success in this study is better explained by male prerut rank and the proportion of days they fight at least twice during the rut.

Lower-ranked males are likely to fight more against other males close in rank distance than against males high above them in the social hierarchy. Lowest-ranked males fought at least twice per day less often compared to the rest of males. In addition, the portion of lowest-ranked males seemed to win fights more often than those ranked in the middle. We suspect that lower-ranked males might engage in more fights with other closely-ranked males, thus resulting in higher proportion of wins than normally expected for subordinate males. Fallow deer males are likely to fight others that are near in rank distance (Bartoš et al., 2007), given that they have a greater chance of winning (Tibbetts et al., 2022). Lower-ranked males, many of which are probably still young (4-5 years old), potentially avoid fights against higher-ranked males, and thus reduce the probability of injury. Additionally, fights among lower-ranked males might help them to gain experience in contests. Indeed, in pigs (*Sus scrofa*) and fallow deer, experienced males assess each other for longer before engaging in a fight (Camerlink et al., 2017; McElligott et al., 1998; Oldham et al., 2020).

We found that higher-ranked males secured most matings throughout the rut, while some lower-ranked males secured some matings towards the end. Higher-ranked males fought at least twice per day on a higher proportion of days at the start and peak of the rut compared to lower-ranked males, but not at the end. The daily investment in fights may affect body condition of higher-ranked males, which in turn affects the quality of their vocalizations (e.g. changes in how their calls sound). Some individuals may start switching to self-maintenance behaviors, reducing overall investment in intra-sexual competition towards the end of the rut (Pitcher et al., 2014; Vannoni & McElligott, 2009; Wyman et al., 2021). By contrast, lower-ranked males are involved in more than one fight less often during the rut, and some of them may increase their effort later in the mating season (Mason et al., 2012; Nieminen et al., 2016). This suggests that lower-ranked males successfully use alternative mating tactics in scenarios when fewer higher-ranked males are preoccupied in further disputing dominance relationships.

## CONCLUSION

To conclude, our results indicate that higher-ranked fallow deer males fight more, while maintaining higher mating success than other socially mature males. Our results suggest that high numbers of fights in a day negatively affect mating success the same day, but can result in delayed benefits by increasing mating success the next day. Furthermore, changes in the proportion of days that males fight at least twice may allow lower-ranked males to secure a mating towards the end of the rut. Overall, our results suggest that fighting do not provide immediate but delayed and longer-term fitness benefits (Dehnen et al., 2022a; Tibbetts et al., 2022).

## DATA AVAILABILITY

The datasets analyzed during the current study are available at https://docs.google.com/spreadsheets/d/1rC2zNQUFAhjiVYFzJcML13kflu81ln9c/edit?usp=sharing&ouid=105937763265807551314&rtpof=true&sd=true

## ETHICAL DECLARATIONS

The research was part of a long-term study on the behavior and ecology of a city park fallow deer herd (Briefer et al., 2010, 2013; Farrell et al., 2011; McElligott et al., 2002). Given that the animals were observed in a public park, and no experiments were carried out, no ethical approval was required to conduct the research at the time (1994 and 1995).

Phoenix Park is located close to the center of Dublin, surrounded by the city, and is accessible to the public. Thus, the deer are habituated to human presence and are not usually disturbed by people standing nearby. Behavioral data collection was usually carried out at a minimum distance of about 30 m from the animals, using spotting scopes mounted on tripods, and did not cause disturbance to their normal activities.

## REFERENCES

Apollonio, M, Di Vittorio, I (2004) Feeding and reproductive behaviour in fallow bucks (Dama dama). Naturwissenschaften 91:579–584. https://doi.org/10.1007/s00114-004-0574-0

Apollonio, M, Festa-Bianchet, M, & Mari, F (1989) Correlates of copulatory success in a fallow deer lek. Behav Ecol Sociobiol 25:89–97. https://doi.org/10.1007/bf00302925

Bartoš, L, FriČová, B, Bartošová-Víchová, J, Panamá, J, Šustr, P, & Šmídová, E (2006) Estimation of the probability of fighting in fallow deer (Dama dama) during the rut. Aggressive Behav 33:7–13. https://doi.org/10.1002/ab.20162

Bates, D, Mächler, M, Bolker, B, & Walker, S (2015) Fitting Linear Mixed-Effects Models Using lme4. J Stat Softw 67:1–48. https://doi.org/10.18637/jss.v067.i01

Bowyer, R T, McCullough, D R, Rachlow, J L, Ciuti, S, & Whiting, J C (2020) Evolution of ungulate mating systems: Integrating social and environmental factors. Ecol Evol 10:5160–5178. https://doi.org/10.1002/ece3.6246

Briefer, E, Farrell, M E, Hayden, T J, & McElligott, A G (2013) Fallow deer polyandry is related to fertilization insurance. Behav Ecol Sociobiol 67:657–665. https://doi.org/10.1007/s00265-013-1485-x

Briefer, E, Vannoni, E, & McElligott, A G (2010) Quality prevails over identity in the sexually selected vocalisations of an ageing mammal. BMC Biol 8. https://doi.org/10.1186/1741-7007-8-35

Burnham, K P, Anderson, D R, & Huyvaert, K P (2010) AIC model selection and Multimodel Inference in behavioral ecology: Some background, observations, and comparisons. Behav Ecol Sociobiol 65:23–35. https://doi.org/10.1007/s00265-010-1029-6

Camerlink, I, Turner, S P, Farish, M, & Arnott, G (2017) The influence of experience on contest assessment strategies. Sci Rep-UK 7. https://doi.org/10.1038/s41598-017-15144-8

Chase, I D, Coelho, D, Lee, W, Mueller, K, & Curley, J P (2022) Networks never rest: An investigation of network evolution in three species of animals. Soc Networks 68:356–373. https://doi.org/10.1016/j.socnet.2021.09.002

Ciuti, S, & Apollonio, M (2016) Reproductive timing in a lekking mammal: Male fallow deer getting ready for female estrus. Behav Ecol 27:1522–1532. https://doi.org/10.1093/beheco/arw076

Clutton-Brock, T H, & Huchard, E (2013) Social competition and selection in males and females. Philos T R Soc B 368:20130074. https://doi.org/10.1098/rstb.2013.0074

Clutton-Brock, T H, Albon, S D, Gibson, R M, & Guinness, F E (1979) The Logical Stag: Adaptive aspects of fighting in red deer (Cervus elaphus L.). Anim Behav 27:211–225. https://doi.org/10.1016/0003-3472(79)90141-6

Clutton-Brock, T H, Green, D, Hiraiwa-Hasegawa, M, & Albon, S D (1988) Passing the buck: Resource defence, lek breeding and mate choice in fallow deer. Behav Ecol Sociobiol 23:281–296. https://doi.org/10.1007/bf00300575

David, H A (1987) Ranking from unbalanced paired-comparison data. Biometrika 74:432–436. https://doi.org/10.1093/biomet/74.2.432

Dehnen, T, Arbon, J J, Farine, D R, & Boogert, N J (2022) How feedback and feed-forward mechanisms link determinants of social dominance. Biol Rev. https://doi.org/10.1111/brv.12838

Dehnen, T, Papageorgiou, D, Nyaguthii, B, Cherono, W, Penndorf, J, Boogert, N J, & Farine, D R (2022) Costs dictate strategic investment in dominance interactions. Philos T R Soc B 377. https://doi.org/10.1098/rstb.2020.0447

Ellis, L (1995) Dominance and reproductive success among nonhuman animals: A cross-species comparison. Ethol Sociobiol 16:257–333. https://doi.org/10.1016/0162-3095(95)00050-u

Emery Thompson, M. & Georgiev, A V (2014) The high price of success: Costs of mating effort in male primates. Int J Primatol 35:609–627. https://doi.org/10.1007/s10764-014-9790-4

Emlen, S & Oring, L (1977) Ecology, sexual selection, and the evolution of mating systems. Science 197:215-223. DOI: 10.1126/science.327542

Fanjul, M S, & Zenuto, R R (2017) Female choice, male dominance and condition-related traits in the polygynous subterranean rodent Ctenomys talarum. Behav Process 142:46–55. https://doi.org/10.1016/j.beproc.2017.05.019

Farrell, M E, Briefer, E, & McElligott, A G (2011) Assortative mating in fallow deer reduces the strength of sexual selection. PLoS ONE 6. https://doi.org/10.1371/journal.pone.0018533

Foley, A M, Hewitt, D G, DeYoung, R W, Schnupp, M J, Hellickson, M W, & Lockwood, M A (2018) Reproductive effort and success of males in scramble-competition polygyny: Evidence for trade-offs between foraging and mate search. J Anim Ecol 87:1600–1614. https://doi.org/10.1111/1365-2656.12893

Gammell, M P, de Vries, H, Jennings, D J, Carlin Caitríona M, & Hayden, T J (2003) David’s score: A more appropriate dominance ranking method than Clutton-Brock et al.’s index. Anim Behav 66:601–605. https://doi.org/10.1006/anbe.2003.2226

Gerber, L R, González-Suárez, M, Hernández-Camacho, C J, Young, J K, & Sabo, J L (2010) The cost of male aggression and polygyny in California Sea Lions (Zalophus californianus). PLoS ONE 5. https://doi.org/10.1371/journal.pone.0012230

Halekoh, U, & Højsgaard, S (2014) A Kenward-Roger Approximation and Parametric Bootstrap Methods for Tests in Linear Mixed Models – The R Package pbkrtest. J Stat Softw 59:1–32. https://doi.org/10.18637/jss.v059.i09

Hartig, F (2020) DHARMa: Residual Diagnostics for Hierarchical (Multi-Level / Mixed) Regression Models. R package version 0.3.3.0. https://CRAN.R-project.org/package=DHARMa

Hobson, E A, Mønster, D, & DeDeo, S (2021) Aggression heuristics underlie animal dominance hierarchies and provide evidence of group-level social information. Proc Natl Acad Sci USA 118. https://doi.org/10.1073/pnas.2022912118

House, C M, Rapkin, J, Hunt, J, & Hosken, D J (2019) Operational sex ratio and density predict the potential for sexual selection in the broad-horned beetle. Anim Behav 152:63–69. https://doi.org/10.1016/j.anbehav.2019.03.019

Hsu, Y, & Wolf, L L (2001) The winner and loser effect: What fighting behaviours are influenced? Anim Behav 61:777–786. https://doi.org/10.1006/anbe.2000.1650

Lewis, R J (2022) Aggression, rank and power: Why hens (and other animals) do not always peck according to their strength. Philos T R Soc B 377. https://doi.org/10.1098/rstb.2020.0434

Lukas, D, & Clutton-Brock, T (2014) Costs of mating competition limit male lifetime breeding success in polygynous mammals. Proc R Soc B Biol Sci 281:20140418. https://doi.org/10.1098/rspb.2014.0418

Magaña, M, Alonso, J C, & Palacín, C (2011) Age-related dominance helps reduce male aggressiveness in great bustard leks. Anim Behav 82:203–211. https://doi.org/10.1016/j.anbehav.2011.04.014

Mason, T H, Stephens, P A, Willis, S G, Chirichella, R, Apollonio, M, & Richards, S A (2012) Intraseasonal variation in reproductive effort: Young males finish last. Am Nat 180:823–830. https://doi.org/10.1086/668082

McElligott, A G, & Hayden, T J (2000) Lifetime mating success, sexual selection and life history of fallow bucks (Dama dama). Behav Ecol Sociobiol 48:203–210. https://doi.org/10.1007/s002650000234

McElligott, A G, Altwegg, R, & Hayden, T J (2002) Age-specific survival and reproductive probabilities: Evidence for senescence in male fallow deer (Dama dama). P Roy Soc Lond B Bio 269:1129–1137. https://doi.org/10.1098/rspb.2002.1993

McElligott, A G, Gammell, M P, Harty, H C, Paini, D R, Murphy, D T, Walsh, J T, & Hayden, T J (2001) Sexual size dimorphism in fallow deer (Dama dama): Do larger, heavier males gain greater mating success? Behav Ecol Sociobiol 49:266–272. https://doi.org/10.1007/s002650000293

McElligott, A G, Mattiangeli, V, Mattiello, S, Verga, M, Reynolds, C A, & Hayden, T J (1998) Fighting tactics of fallow bucks (Dama dama, Cervidae): Reducing the risks of serious conflict. Ethology 104:789–803. https://doi.org/10.1111/j.1439-0310.1998.tb00112.x

McElligott, A G, O’Neill, K P, & Hayden, T J (1999) Cumulative long-term investment in vocalization and mating success of Fallow Bucks, Dama dama. Anim Behav 57:1159–1167. https://doi.org/10.1006/anbe.1999.1076

McElligott, AG, Naulty, F, Clarke, WV & Hayden, T J (2003) The somatic cost of reproduction: what determines reproductive effort in prime-aged fallow bucks? Evol Ecol Res 5:1239–1250. https://doi.org/10.5167/uzh-402

Moore, N P, Kelly, P F, Cahill, J P, & Hayden, T J (1995) Mating strategies and mating success of Fallow (Dama dama) bucks in a non-lekking population. Behav Ecol Sociobiol 36:91–100. https://doi.org/10.1007/bf00170713

Mysterud, A, Bonenfant, C, Loe, L E, Langvatn, R, Yoccoz, N G, & Stenseth, N C (2008) The timing of male reproductive effort relative to female ovulation in a capital breeder. J Anim Ecol 77:469–477. https://doi.org/10.1111/j.1365-2656.2008.01365.x

Naulty, F, Harty, H, & Hayden, T J (2013) Freedom to choose: Unconstrained mate-searching behaviour by female fallow deer (Dama dama). Folia Zool 62:143–154. https://doi.org/10.25225/fozo.v62.i2.a10.2013

Nieminen, E, Kervinen, M, Lebigre, C, & Soulsbury, C D (2016) Flexible timing of reproductive effort as an alternative mating tactic in black grouse (Lyrurus tetrix) males. Behaviour 153:927–946. https://doi.org/10.1163/1568539x-00003374

Oldham, L, Camerlink, I, Arnott, G, Doeschl-Wilson, A, Farish, M, & Turner, S P (2020) Winner–loser effects overrule aggressiveness during the early stages of contests between pigs. Sci Rep-UK 10. https://doi.org/10.1038/s41598-020-69664-x

Pitcher, B J, Briefer, E F, Vannoni, E, & McElligott, A G (2014) Fallow bucks attend to vocal cues of motivation and fatigue. Behav Ecol 25:392–401. https://doi.org/10.1093/beheco/art131

Shizuka, D, & McDonald, D B (2012) A social network perspective on measurements of dominance hierarchies. Anim Behav 83:925–934. https://doi.org/10.1016/j.anbehav.2012.01.011

Silk, M J, Cant, M A, Cafazzo, S, Natoli, E, & McDonald, R A (2019) Elevated aggression is associated with uncertainty in a network of dog dominance interactions. Proc R Soc B Biol Sci 286:20190536. https://doi.org/10.1098/rspb.2019.0536

Tibbetts, E A, Pardo-Sanchez, J, & Weise, C (2022) The establishment and maintenance of dominance hierarchies. Philos T R Soc B 377. https://doi.org/10.1098/rstb.2020.0450

Vannoni, E, & McElligott, A G (2009) Fallow bucks get hoarse: Vocal fatigue as a possible signal to conspecifics. Anim Behav 78:3–10. https://doi.org/10.1016/j.anbehav.2009.03.015

Willisch, C S, & Neuhaus, P (2010) Social dominance and conflict reduction in rutting male alpine ibex, Capra ibex. Behav Ecol 21:372–380. https://doi.org/10.1093/beheco/arp200

Willisch, C S, Biebach, I, Koller, U, Bucher, T, Marreros, N, Ryser-Degiorgis, M-P, Keller, L F, & Neuhaus, P (2011) Male reproductive pattern in a polygynous ungulate with a slow life-history: The role of age, social status and alternative mating tactics. Evol Ecol 26:187–206. https://doi.org/10.1007/s10682-011-9486-6

Wyman, M T, Pinter-Wollman, N, & Mooring, M S (2021) Trade-offs between fighting and breeding: A social network analysis of bison male interactions. J Mammal 102:504–519. https://doi.org/10.1093/jmammal/gyaa172

