## Supplementary Information for "Delayed benefits for fallow bucks: more fights decrease same day mating success, but may increase matings the next day"

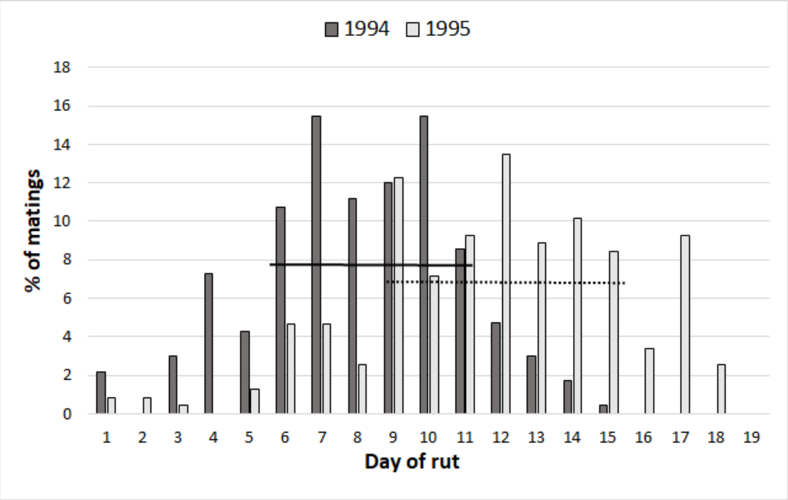


**Fig. S1** Percentage of matings per day of rut in 1994 and 1995. Solid (1994) and dashed (1995) horizontal lines show the days that were attributed to the peak of the rut, depending on the minimum percentage of matings for a day (7.7% in 1994 and 6.8% in 1995)

**Table S1** Relationship between rank (prerut hierarchical dominance) and 1) mean fight rate (calculated per buck per day), 2) mean mating rate (calculated per buck per day), and 3) mean outcome of fights (1 = win; 0 = loss). For each relationship, three different models (factor of interest fitted as a linear, quadratic or logarithmic term) were tested against the model without the factor of interest. (K = number of parameters; AICc = Akaike Information Criterion with a correction for small sample sizes; DAICc = delta of AICc against value of model with lowest AICc)

| | | **Fight rate[1]** | | **Mating rate[2]** | | **Outcome[3]** | |
|---|---|---|---|---|---|---|---|
| | **K** | AICc | DAICc | AICc | DAICc | AICc | DAICc |
| **No factor of interest** | **6** | 1525.5 | 27.7 | 733.19 | 59.99 | 1272.6 | 22.6 |
| **Linear** | **7** | 1501 | 3.2 | 682.39 | 9.19 | 1263.4 | 13.4 |
| **Quadratic** | **8** | 1497.8 | 0 | 673.2 | 0 | 1250 | 0 |
| **Logarithmic** | **7** | 1512.1 | 14.3 | 674.65 | 1.45 | 1253.1 | 3.1 |

**Table S2** Relationship between mean fight rate (calculated per buck per day) and 1) mean mating rate that day (calculated per buck per day), and 2) mean mating rate the next day (calculated per buck per day). For each relationship, three different models (factor of interest fitted as a linear, quadratic or logarithmic term) were tested against the model without the factor of interest. (K = number of parameters; AICc = Akaike Information Criterion with a correction for small sample sizes; DAICc = delta of AICc against value of model with lowest AICc)

| | | **Mating rate[1]** | | **Mating rate the next day[2]** | |
|---|---|---|---|---|---|
| | **K** | AICc | DAICc | AICc | DAICc |
| **No factor of interest** | **6** | 676.53 | 23.86 | 651.57 | 12.87 |
| **Linear** | **7** | 665.15 | 12.48 | 641.52 | 2.82 |
| **Quadratic** | **8** | 652.67 | 0 | 640.9 | 2.2 |
| **Logarithmic** | **7** | 656.99 | 4.32 | 638.7 | 0 |

**Table S3** Relationship between mean outcome of fights (1 = win; 0 = loss) and 1) mean mating rate that day (calculated per buck per day), and 2) mean mating rate the next day (calculated per buck per day). For each relationship, three different models (factor of interest fitted as a linear, quadratic or logarithmic term) were tested against the model without the factor of interest. (K = number of parameters; AICc = Akaike Information Criterion with a correction for small sample sizes; DAICc = delta of AICc against value of model with lowest AICc)

| | | **Mating rate[1]** | | **Mating rate the next day[2]** | |
|---|---|---|---|---|---|
| | **K** | AICc | DAICc | AICc | DAICc |
| **No factor of interest** | **6** | 322.77 | 0 | 135.01 | 0 |
| **Linear** | **7** | 323.45 | 0.68 | 135.96 | 0.95 |
| **Quadratic** | **8** | 323.45 | 0.68 | 135.96 | 0.95 |
| **Logarithmic** | **7** | 323.45 | 0.68 | 135.96 | 0.95 |

**Table S4** Relationship between rank (prerut hierarchical dominance) and mean fight rate (calculated per buck per day) at the Start, Peak and End of the rut. For each period, three different models (factor of interest fitted as a linear, quadratic or logarithmic term) were tested against the model without the factor of interest. (K = number of parameters; AICc = Akaike Information Criterion with a correction for small sample sizes; DAICc = delta of AICc against value of model with lowest AICc)

| **FIGHTS** | | **Start** | | **Peak** | | **End** | |
|---|---|---|---|---|---|---|---|
| | K | AICc | DAICc | AICc | DAICc | AICc | DAICc |
| **No factor of interest** | 6 | 595.25 | 6.4 | 663.27 | 22.02 | 285.91 | 1.24 |
| **Linear** | 7 | 592.83 | 3.98 | 641.25 | 0 | 285.05 | 0.38 |
| **Quadratic** | 8 | 588.85 | 0 | 642.87 | 1.62 | 284.67 | 0 |
| **Logarithmic** | 7 | 596.33 | 7.48 | 645.08 | 3.83 | 286.7 | 2.03 |

**Table S5** Relationship between rank (prerut hierarchical dominance) and mean mating rate (calculated per buck per day) at the Start, Peak and End of the rut. For each period, three different models (factor of interest fitted as a linear, quadratic or logarithmic term) were tested against the model without the factor of interest. (K = number of parameters; AICc = Akaike Information Criterion with a correction for small sample sizes; DAICc = delta of AICc against value of model with lowest AICc)

| **MATINGS** | | **Start** | | **Peak** | | **End** | |
|---|---|---|---|---|---|---|---|
| | K | AICc | DAICc | AICc | DAICc | AICc | DAICc |
| **No factor of interest** | 6 | 196.92 | 25.98 | 424.86 | 42.18 | 142.89 | 5.69 |
| **Linear** | 7 | 171.89 | 0.95 | 387.09 | 4.41 | 141.21 | 4.01 |
| **Quadratic** | 8 | 170.94 | 0 | 382.68 | 0 | 139.44 | 2.24 |
| **Logarithmic** | 7 | 171.16 | 0.22 | 383.63 | 0.95 | 137.2 | 0 |
